# Thermal regime during parental sexual maturation, but not during offspring rearing, modulates DNA methylation in brook charr (*Salvelinus fontinalis*)

**DOI:** 10.1101/2022.02.25.481661

**Authors:** Clare J Venney, Kyle W Wellband, Eric Normandeau, Carolyne Houle, Dany Garant, Céline Audet, Louis Bernatchez

## Abstract

Epigenetic inheritance can result in plastic responses to changing environments being faithfully transmitted to offspring. However, it remains unclear how epigenetic mechanisms such as DNA methylation can contribute to multigenerational acclimation and adaptation to environmental stressors. Brook charr (*Salvelinus fontinalis*), an economically important salmonid, is highly sensitive to thermal stress, and is of conservation concern in the context of climate change. We studied the effects of temperature during parental sexual maturation and offspring rearing on whole-genome DNA methylation in brook charr juveniles (fry). Parents were split between warm and cold temperatures during sexual maturation, mated in controlled breeding designs, then offspring from each family were split between warm (8°C) and cold (5°C) rearing environments. We found 188 differentially methylated regions (DMRs) due to parental maturation temperature after controlling for family structure. In contrast, offspring rearing temperature had a negligible effect on offspring methylation. Stable intergenerational inheritance of DNA methylation and minimal plasticity in progeny could result in transmission of acclimatory epigenetic states to offspring, priming them for a warming environment. Our findings have implications pertaining to the role of intergenerational epigenetic inheritance in response to ongoing climate change.

## Introduction

Climate change is a pervasive threat to global biodiversity and is expected to have profound effects on the resilience and abundance of species [1]. In Canada, the mean annual temperature has increased by 1.7°C from 1948 to 2016 and is expected to increase over the next 30 years in both low and high emission scenarios [2]. As sea surface temperatures increase, it is expected that biomass of aquatic organisms such as fish will decrease, resulting in considerable economic losses [3]. With most of Eastern Canada experiencing moderate increases in mean annual temperatures and temperature extremes, fish catch is predicted to decrease and considerable declines in fish stocks are forecast due to long-term increases in temperature [3]. Therefore, a thorough understanding of the evolutionary mechanisms through which fishes can respond to climate change is a priority for the conservation of fish stocks [4–6].

There is ample evidence for plasticity to thermal stress in fish, as observed through differences in physiology [7,8], morphology [8,9], and behaviour [9,10], among other traits. These plastic responses are often due to underlying differences in gene expression driven by thermal regimes [11–15]. In some instances, intergenerational plasticity can occur wherein plastic phenotypes are passed on from parent to offspring [16–18]. Thus, parents have the potential to pass on phenotypes to offspring based on environmental experience, which can influence offspring fitness [19,20] and fitness-related traits [17,21]. The underlying mechanisms for intergenerational plasticity have not been thoroughly characterized but may be partially due to epigenetic inheritance.

Epigenetic inheritance refers to the transmission of epigenetic marks (e.g., DNA methylation) from parent to progeny that can modify offspring gene expression, phenotype, and fitness [22,23]. DNA methylation is an important regulator of transcription and has been shown to change in response to temperature [24–27], differences in rearing environment [28,29], and other factors. Previous studies have shown that epigenetic changes are passed on to offspring due to parental exposure to factors such as thermal stress [30–35] and hatchery rearing [36–38]. Thus, epigenetic inheritance is common in response to parental experiences and can have important implications for offspring phenotype and fitness [22,23]. In particular, epigenetic inheritance could help future generations by preparing them for warming climates, and thus merits further study as a mechanism for species to cope with climate change and increasingly inhospitable environments [35].

Brook charr (*Salvelinus fontinalis*) is one of the most prized sport fish in Eastern Canada [39]. The brook charr sport fishery supports 3 000 jobs and brings in $340 million in revenue each year in Québec alone [40]. Large-scale stocking occurs each year to replenish stocks and preserve the sport fishery, costing a total of $5 million yearly [41]. Nearly 650 000 kg of brook charr, representing ∼70% of all stocked fish, are released into Québec lakes each year [41]. Brook charr is highly sensitive to thermal stress [39,42,43] and thus to climate change. An increase in summer air temperature of 1°C is sufficient to delay spawning by one week and reduce redd production by 22 to 65% [42]. Populations show little variation in their capacity to respond to temperature changes [44,45], though there is some evidence that populations from cooler climates have lower thermal tolerance [46]. Based on predicted climate scenarios, it is expected that suitable brook charr habitat will decrease over time across much of its native range [43,47–50], including in Québec and Eastern Canada where suitable habitat would shift to the northeast [51], resulting in reduced growth and survival of individuals and ultimately persistence of populations [43,45,52]. Due to the thermal sensitivity and economic importance of the species, a thorough understanding of the mechanisms through which brook charr acclimate and cope with thermal stress is needed to refine predictions of climate change impacts on this species.

An important unresolved question on epigenetic inheritance is whether offspring environmental influences supersede environmentally induced epigenetic states inherited from their parents. If inherited methylation patterns persist regardless of offspring environment, this would allow the persistence of epigenetic states that could prime offspring for warming environments and provide additional adaptive capacity to populations experiencing warming. However, high offspring plasticity in response to their own environment may reduce the advantage of epigenetic inheritance if offspring rapidly and appropriately acclimate to their perceived environment, overriding inherited epigenetic marks. Here we use a reciprocally crossed design of parents and offspring reared under contemporary and warming conditions combined with whole-epigenome sequencing to assess the relative importance of parental and offspring experience on the offspring epigenome. The results of this study add to our knowledge of the molecular mechanisms through which organisms can respond to climate change, furthering our understanding of the role that epigenetic inheritance plays in acclimation and evolutionary inheritance.

## Methods

### Fish rearing and breeding design

Brook charr from the Laval strain [53], a captive strain descended from the Laval River in Québec and reared for six generations, were used as broodstock for the experiment. The fish were held at the ISMER (Institute de Sciences de la Mer de Rimouski) wet lab facilities at Université du Québec à Rimouski and were split between two thermal regimes shortly before sexual maturation: warm and cold treatments, separated by ∼2°C. After sexual maturation, 2×2 breeding crosses were created when possible (see Figure S1 for breeding design). Eggs from each family were split in two batches and sent to the LARSA (Laboratoire de Recherche en Sciences Aquatiques) at Université Laval. There, half of the eggs from each family were incubated at a warm 8°C thermal regime treatment and the other half at 5°C which was representative of end of fall water temperature. Both temperature treatments were maintained through egg development and the yolk-sac fry period. This design allowed us to determine the relative influence of parental vs. offspring rearing temperature on the offspring epigenome. Upon yolk sac resorption, the offspring from both treatments were held at ∼8.5°C, a typical rearing temperature for this developmental stage, to encourage feeding and minimize mortality. Fry from each family were sampled after two months of exogenous feeding at an approximate size of 85 mm/5 g. All fish were humanely euthanized with an overdose solution of tricaine methanesulfonate (200 ppm). Liver tissues were immediately dissected and preserved in RNAlater for future analysis. All animal care was performed humanely under permit VRR-18-111.

### Parentage analysis

Parentage analysis was performed to confirm that the sampled fish corresponded to the correct family, as described in [54]. Briefly, DNA was extracted from fin clips from parents and offspring, amplified at 12 microsatellite loci [55], and visualized on an AB3500 automated DNA sequencer. GeneMapper V6 (Applied Biosystem) was used to determine allele lengths, which were imported into both Cervus v3.0.7 [56] and COLONY v2.0.6 [57]. Parentage analysis was performed using COLONY’s full likelihood approach and Cervus’ 90% confidence likelihood approach. Parentage assignment was considered successful when both programs identified the same set of parents. If the identified parents were not crossed in the breeding design, the next most probable parentage assignment was used. If both programs did not suggest the same pair, the most probable pair between the possible crosses was assigned. Inability to assign both parents to an individual resulted in a failed assignation.

### DNA extraction and whole-genome bisulfite sequencing

Livers were selected for 54 male offspring: two offspring per family, except for two families where only one offspring had confirmed parentage and sex (Figure S1). Liver tissue was used due to its homogeneity in cell types and involvement in metabolism and growth [58]. For each combination of parental and offspring temperatures (e.g., warm parental maturation and warm offspring maturation), one 2×2 cross and one partial 2×2 missing a family was used (Figure S1). DNA was extracted using a salt-based extraction protocol, quantified, and checked for quality. Offspring sex was verified using a genetic sex marker for salmonids [59] and only male offspring were used for sequencing to eliminate sex-specific methylation effects. Library preparation, quality control, and sequencing were performed by the Centre d’expertise et de services of Génome Québec, Montréal, Canada. Methyl-Seq with approximately 15X coverage was performed on the Illumina NovaSeq6000 using S4 flow cells for 150 bp paired-end reads across four sequencing lanes.

### Sequence data processing

Data were trimmed using fastp [60] to remove sequences under 100 bp and with phred scores below 25, and the first and last nucleotides were trimmed. Bwa-meth (https://github.com/brentp/bwa-meth) was used to align the sequence data to the lake trout (*S. namaycush*) genome (SaNama v1.0; NCBI Refseq: GCF_016432855.1) [61], a closely related sister species of brook charr [62]. Duplicate reads were removed from the bam files with picard tools v1.119 *MarkDuplicates* (https://github.com/broadinstitute/picard). MethylDackel’s *mbias* function was used to inform trimming of noisy, biased regions at the beginnings and ends of reads (https://github.com/dpryan79/MethylDackel). Methylation was called using MethylDackel *extract* and the paired end reads were merged to produce bedGraph and methylKit files. The pipeline is available at https://github.com/enormandeau/bwa-meth_pipeline.

### Identifying and masking SNPs from methylation data

Existing SNP data from pooled sire DNA including the eight sires from this study and 32 other males generated by a related study (Wellband et al. in prep) was used to identify and mask C/T SNPs which cannot be differentiated from true methylation calls in the Methyl-Seq data. SNP data were trimmed with fastp using the same quality requirements as the methylation data (minimum length of 100 bp, minimum phred score of 25, and trimming first and last bases). Sequences were aligned to the lake trout genome using bwa [63], duplicate reads were removed using picard tools *MarkDuplicates*, and overlapping reads were clipped using bamUtil *clipOverlap* [64]. Freebayes [65] was used to call SNPs covered by between 10 and 100 reads with a minimum allele frequency of 0.01 and at least two reads for the alternative allele. A list of C/T and A/G SNPs was extracted and exported into bed format. The pipeline is available online at https://github.com/kylewellband/CT-poly-wgbs.

### DNA methylation analysis and jackknifing

SNPs were masked from the methylation data (bedGraph and methylKit files) using bedtools *intersect* with the *-v* option [66]. The bedGraph files were read into bsseq [67] and filtered to require between five and 80 reads per CpG in at least 80% of the samples (i.e., 44 of 54 individuals). Methylation data were smoothed over 500 bp regions using the built-in moving average algorithm in DSS to control for spatial correlation of methylation levels among proximal CpGs [68]. A beta-binomial generalized linear model for the effects of adult temperature, offspring temperature, and their interaction was implemented in DSS to identify differentially methylated loci (DMLs). DMLs were considered significant if they had a p-value of less than 0.001 and differentially methylated regions (DMRs) were then called based on the DML results for each term in the model using DSS.

We used jackknife resampling to confirm that DMRs were not driven by family effects, which we were not able to directly control for in DSS due to model overfitting. Based on the hierarchical clustering for the full dataset, we created 14 data subsets in bsseq, each with all offspring from a given full-sibling family dropped from the analysis. DML and DMR detection were performed with DSS for each subset with a p-value cut-off of 0.05 to allow for some variation in the significance of DMLs in subsets. The DMR result files were converted to bed format and bedtools *intersect* was used to determine which subsets had DMRs that overlapped with the DMRs of the full dataset. Subset DMRs had to overlap at least 80% of the length of the original DMR to be considered equivalent. Subset DMRs obtained from jackknife resampling that satisfied this condition in all subsets were considered verified DMRs. All codes are available at https://github.com/cvenney/methylUtil. Results were visualized using the R package ComplexHeatmap [69].

### Annotation and gene ontology enrichment analysis

Gene ontology analysis was performed for the jackknife verified DMRs for adult maturation temperature. First, the GCF_016432855.1_SaNama_1.0 genome and transcriptome available from GenBank were used with the GAWN v0.3.5 pipeline (github.com/enormandeau/gawn), using the default parameters, to annotate the transcripts and find the DMRs (i) directly overlapping transcripts, and (ii) within ±5 kb of transcripts. Using the lists of DMRs and annotated transcripts, GO enrichment analysis was done using the go_enrichment v1.0.0 pipeline (github.com/enormandeau/go_enrichment), using the default parameters. A Benjamini-Hochberg false discovery rate (FDR) correction was used to correct for multiple comparisons with a critical p-value of p<0.05.

### Redundancy analysis for family and offspring temperature effects

We used redundancy analysis (RDA) to determine whether family and offspring temperature affected overall methylation levels in offspring from each adult maturation temperature. The dataset was split by adult temperature to form two datasets which underwent the same analysis. MethylKit files were imported into R and filtered to include only CpG sites with coverage between five and 80 reads using the methylKit package [70]. CpG sites with data for all individuals were united into one large data frame using methylKit, which was transposed and used for further analysis. RDAs for the effects of family and offspring temperature on whole-genome methylation were performed for each dataset in the package vegan [71]. Significance was tested using an ANOVA-like permutation test with 999 permutations in vegan.

## Results

We obtained an average of 352 941 309 ± 50 216 202 raw Methyl-Seq reads per individual, with an average of 148 451 587 ± 35 187 570 alignments to the lake trout genome after all processing and deduplication. We attained an average of 10.5X ± 2.30 coverage across 16 106 361 analyzed CpG sites for each sample.

### Differential methylation analysis and jackknifing

Adult temperature had the greatest influence on offspring DNA methylation: we identified 464 DMRs due to adult sexual maturation temperature, 34 DMRs due to offspring rearing temperature, and 11 DMRs due to an interaction between the two main terms. Hierarchical clustering of DMRs driven by adult temperature resulted in clear differentiation between the methylation patterns of offspring from warm vs. cold acclimated parents (Figure S2). Further clustering of offspring from full-sibling families was evident within groups (Figure S2), thus jackknife resampling was performed by dropping offspring from each family one by one and rerunning the analysis. Jackknifing resulted in the verification of 188 DMRs based on adult maturation temperature (Figure 1) and 10 DMRs due to offspring rearing temperature. No adult x offspring temperature DMRs persisted after jackknifing.

**Figure 1:**
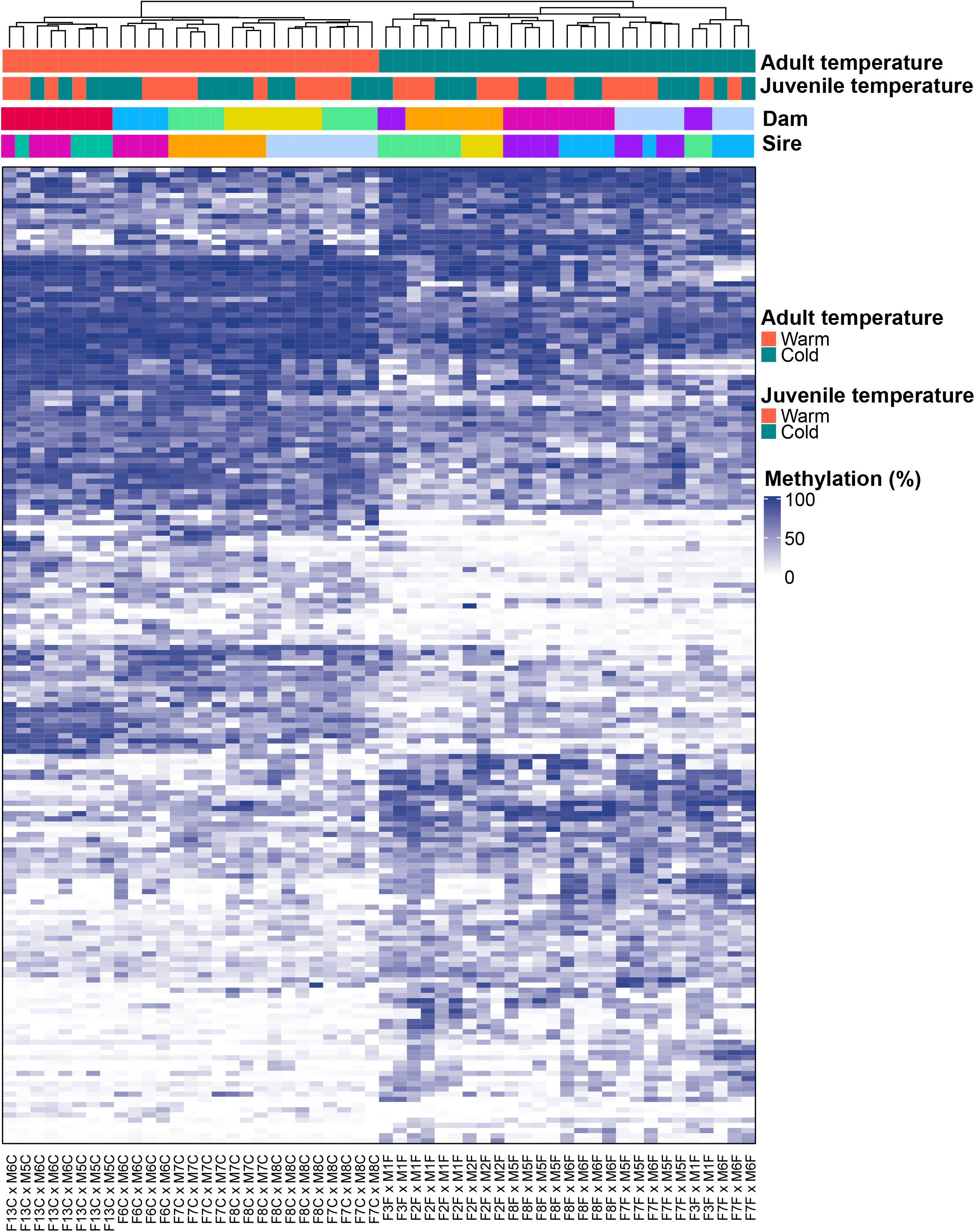
DMR jackknifing confirmed 188 DMRs due to adult sexual maturation temperature. Jackknifing was performed using 14 data subsets, each lacking one full-sibling family. Hierarchical clustering of samples across the x-axis was performed using Euclidean distance and shows strong clustering of samples based on adult maturation temperature. Unique dams and sires are colour-coded on the x-axis at the top of the figure, with consistent blocks of colour indicating clustering based on either maternal or paternal identity. Percent methylation is shown from 0% (white) to 100% (dark blue) in the heatmap with clear differences between offspring from warm and cold-matured parents.

### Functional annotation of DMRs

Fifty-six of the 188 jackknife verified adult temperature DMRs directly overlapped with transcripts. No gene ontology (GO) terms showed significant overrepresentation after FDR correction, either with direct transcript overlaps or ±5 kb from the DMRs. However, many GO terms had multiple transcripts associated with them, including the biological processes of signal transduction, angiogenesis, cell cycle, brain development, and cell differentiation (Table S1).

### RDAs for family and offspring temperature effects

While parental maturation temperature was the main factor driving differences in offspring methylation, we observed family effects on methylation in RDA analyses. RDA models testing for family and offspring temperature effects on whole-genome methylation for both datasets (warm and cold adult maturation temperature) were significant (warm: p = 0.004, adjusted R^2^ = 0.077; cold: p = 0.002, adjusted R^2^ = 0.05). The family effect was significant in both models (warm: p = 0.004; cold: p = 0.002) but offspring temperature only significantly affected methylation of offspring from warm-matured parents (warm: p = 0.041; cold: p = 0.657). We observed grouping of full-sibling families in both temperatures, though this effect was stronger in offspring descended from cold-matured adults (Figure 2). There was some evidence of half-sibling families clustering based on paternal identity in the warm environment (i.e., M7C, M6C), and based on maternal (F8F) and paternal (M1F) identity in the cold environment (Figure 2A-B).

**Figure 2:**
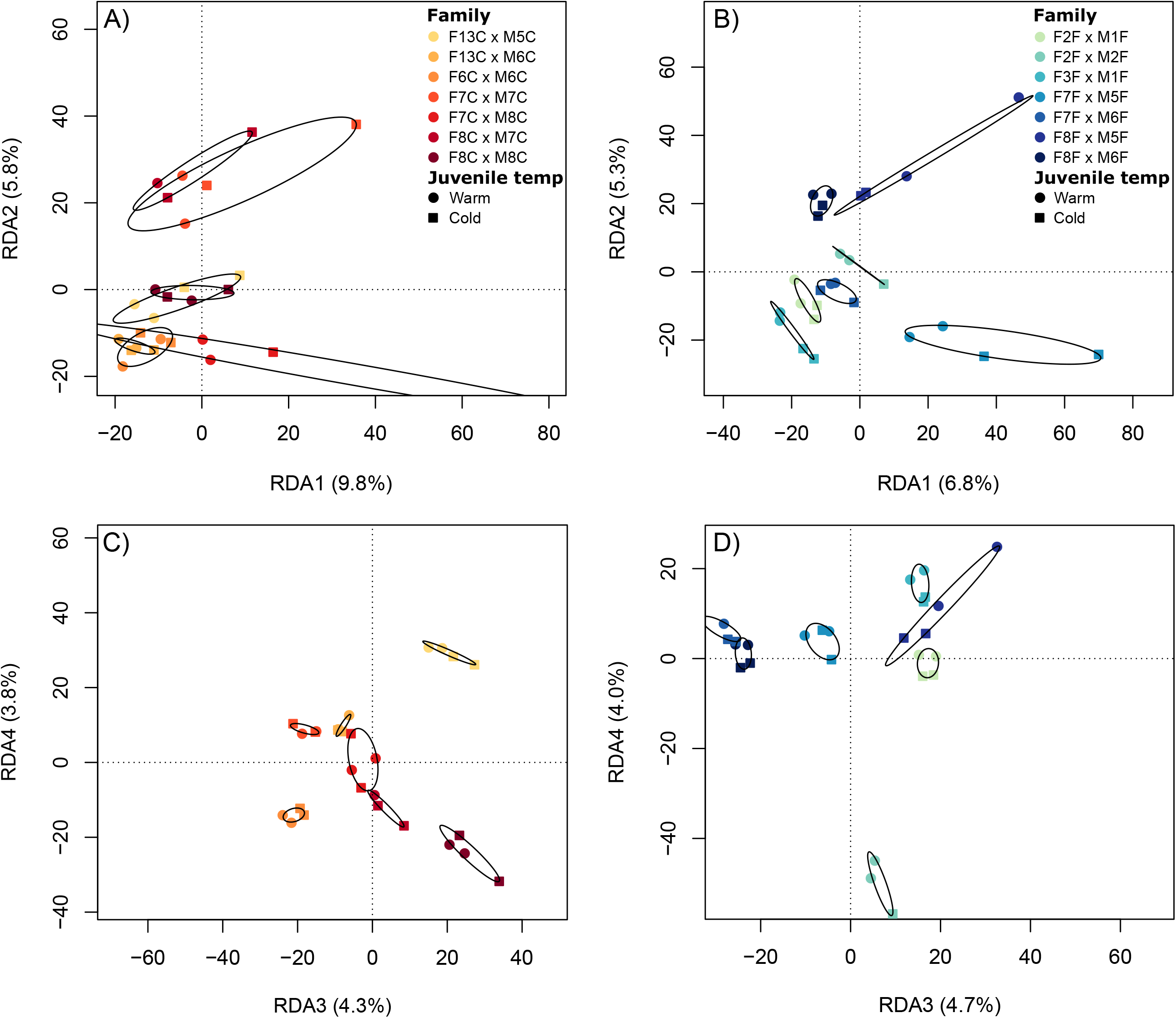
RDAs for family and offspring temperature effects on whole-genome methylation in offspring descended from parents that underwent sexual maturation in warm (8°C, A and C) and cold (5°C, B and D) temperatures. Plots show RDA axes 1 and 2 (A and B) and axes 3 and 4 (C and D). Point colour indicates family of origin while point shape indicates offspring temperature regime.

## Discussion

The main objective of this study was to determine the relative importance of parent versus offspring thermal regimes on offspring DNA methylation. Our results showed that parental sexual maturation temperature, but not offspring rearing temperature, had considerable effects on offspring DNA methylation. This is one of the first studies to assess the relative importance of parental versus offspring rearing environment on offspring DNA methylation, particularly where offspring were split between environments that resembled and differed from the parental environment [22]. Previous studies that employed such reciprocal designs reached contrasting conclusions regarding the relative importance of parental and offspring environment on offspring DNA methylation. Greater parental influences on offspring methylation were reported in the purple sea urchin (*Strongylocentrotus purpuratus*) due to parental temperature and pCO_2_ conditions [72] and in ribwort plantain (*Plantago lanceolata* L.) based on natural multigenerational exposure to varying CO_2_ levels [73]. However, a study in self-fertilizing mangrove rivulus fish (*Kryptolebias marmoratus*) that manipulated parental and offspring environmental enrichment showed that most inherited methylation changes associated with parental environment were lost when offspring were reared in a mismatched environment [74]. A natural reciprocal transplant study in clonally propagated coral *Acropora millepora* found that transplanted corals that altered gene body methylation to resemble local corals had improved fitness-related traits relative to corals that showed minimal plasticity in methylation [75]. Therefore, organisms differ in their ability to override parentally inherited DNA methylation, which could affect the adaptive potential of epigenetic inheritance. Interestingly, the asexually reproducing species (mangrove rivulus and coral) in these studies were more capable of overwriting inherited marks than the sexually reproducing species (purple sea urchin, ribwort plantain, and our study on brook charr), consistent with findings that reproductive mode likely influences the prevalence and persistence of epigenetic inheritance [22]. Since epigenetic variation can persist late into the lifespan of offspring and can be transmitted for multiple generations, inheritance in the absence of offspring plasticity can have profound impacts on phenotype and fitness for generations to come [22,76]. The considerable effects of adult sexual maturation temperature on brook charr offspring methylation reported, coupled with the lack of plasticity in DNA methylation due to offspring temperature, could have significant long-term implications for brook charr populations responding to climate change if offspring cannot override inherited epigenetic marks.

The conclusion that offspring rearing temperature had negligible effects on methylation was unexpected due to overwhelming evidence for thermal regime driving within-generation plastic changes in DNA methylation in fish [24–27,77] and various taxa [35]. It is possible that the 3°C difference between warm and cold rearing temperatures in our study was not sufficient to elicit plasticity in offspring DNA methylation. A lack of sensitivity to slight temperature increases was observed in the spiny chromis damselfish (*Acanthochromis polyacanthus*), which altered oxygen consumption due to a 3°C but not 1.5°C temperature increase [7], and in another fish, the longjaw mudsucker (*Gillichthys mirabilis*), which showed only minor transcriptional differences associated with mild heat stress [78]. For brook charr, this seems unlikely due to high thermal sensitivity [52] and reduction in fitness-related traits due to 1-2°C temperature increases [42,45]. Additionally, the 3°C increase in the warm temperature treatment in our study was sufficient to elicit epigenetic changes in parents that were passed to offspring. It is more likely that the lack of response to offspring temperature is due to an inability of offspring to override inherited DNA methylation, or to detect appropriate temperature cues. Studies have reported variable capacities for plastic responses to the environment through ontogeny [79,80]. Consistent with this, a previous study in European seabass (*Dicentrarchus labrax*) reported altered DNA methylation and gene expression in response to thermal stress in the larval stage, though juvenile free-feeding fish did not show temperature-specific methylation changes [24]. It is also possible that a set of “core” loci respond to both warm and cold temperature treatments through altered DNA methylation, as observed in threespine stickleback (*Gasterosteus aculeatus*) [27]; these “core” loci may not be identified as DMRs in our study as they might respond uniformly to both warm and cold treatments. Overall, we show that adult temperature during sexual maturation has profound effects on offspring methylation regardless of offspring rearing temperature, resulting in stable epigenetic inheritance and low plasticity in response to juvenile brook charr thermal regime.

Based on the maternal match hypothesis, inherited phenotypic differences can be adaptive if offspring environment matches the environment predicted by the parent, but maladaptive if too different [81]. Since research has increasingly identified paternal effects on offspring phenotype across taxa [82], this concept can be expanded to the parental match hypothesis. By the end of the century, average temperature in Canada is expected to increase by 1.8°C under low emission scenarios and by 6.3°C in high emission scenarios [2]. Daily maximum and minimum temperatures are expected to increase by 1.5-6.1°C and 2.8-11.2°C, respectively [2], and extreme temperature events are predicted to gradually increase over the coming years [2,3]. If inherited differences in DNA methylation affect offspring phenotype, epigenetic inheritance due to parental thermal regime could prove adaptive for brook charr due to gradual but predictable warming in Canada, but maladaptive if brook charr are unable to acclimate to and survive transient temperature extremes. The offspring used in this study were sampled at the fry (i.e., early exogenous feeding) stage, thus it is possible that brook charr may exhibit greater plasticity later in development, due to either strong parental effects during early life stages, or developmental canalization resulting in low plasticity during early life [83]. Other studies have identified long-lasting parental effects on gene expression [18] and offspring size [84,85] in brook charr, and a related study using Laval strain brook charr identified persistent parental effects on phenotype past stocking [54]. Persistent parental effects on methylation could result in offspring primed for a warming climate, or epigenetic traps wherein stable epigenetic changes in offspring prove maladaptive but could intensify selection and adaptation to novel environments [86]. Further research into the capacity of offspring to overwrite parentally inherited methylation through ontogeny, and the fitness consequences of heritable epigenetic marks, is needed to determine the permanence and evolutionary consequences of epigenetic inheritance [22].

The family effects observed in this study could be caused by non-genetic parental effects or genetic control of DNA methylation [87], both of which can contribute to epigenetic variation. Due to the consistent grouping of full-sibling families in our analysis, which was stronger for offspring from cold-matured parents, there is likely some extent of genetic control or non-additive effects on DNA methylation. Altered body mass heritability was previously reported due to manipulation of brook charr thermal environment [84], thus it is possible that stressful thermal environments led to increased variation in offspring traits including DNA methylation. Similar increases in offspring variation were reported in dandelion (*Taraxacum* spp.), where parental exposure to salicylic acid increased variation in offspring DNA methylation [88]. The clustering of both maternal and paternal half-sibling families in Figures 1 and 2 suggests that both maternal and paternal effects are acting on methylation, though the clustering was slightly biased towards paternal effects. Early research on epigenetic inheritance in fish suggested that the sperm methylome is primarily inherited while maternal methylation patterns are lost [89,90]. More recently, studies have provided evidence for maternal effects on methylation [91], though the prevalence of maternal effects depends on rearing environment [80]. It is therefore possible that the family effects are influenced by maternal effects, which we were not able to test due to model overfitting (i.e., sample size restrictions), or due to underlying genetic variation driving methylation states. However, it is difficult to disentangle epigenetic and genetic variation [22,76], and thus further research is needed to determine the proximate causes of family effects on methylation.

Our study reinforces the relevance of epigenetic inheritance in response to climate change as epigenetic changes due to parental sexual maturation temperature persisted regardless of offspring thermal environment. Such instances of epigenetic inheritance have the potential to prime offspring for an environment based on parental experience, though they could prove maladaptive for offspring if parental environment is too different from that of the offspring and if the offspring have limited capacity to overwrite inherited methylation. Since climate change will pose a significant threat to brook charr in the coming years [43,47,48,50–52], a thorough understanding of the mechanisms of plasticity through which fish can cope with changing environments is needed [6]. From an applied standpoint, the conservation implications of epigenetic inheritance remain unclear, particularly after release of stocked fish into natural environments. Our study provides a glimmer of hope that brook charr management programs could influence offspring through stable epigenetic changes due to short-term manipulation of parental environment before spawning. Further research into the stability and fitness consequences of epigenetic inheritance is needed to understand the evolutionary implications of epigenetic variation [22]. If future studies prove these instances of epigenetic inheritance to be stable and adaptive, our findings could have significant implications for predicting the survival and persistence of stocked brook charr in warming climates. Our study reinforces the relevance of epigenetic inheritance in intergenerational responses to changes in thermal regime, with the potential to pre-emptively prepare organisms for changing environments. Given the ongoing climate crisis and habitat changes worldwide, a greater understanding of epigenetic and non-genetic heritable sources of variation is critical to understanding the evolutionary potential of organisms.

## Acknowledgements

We thank Gabriel Piette-Lauzière for assistance with sample preparation, Charles Babin for assistance with sampling, and the staff of LARSA (Université Laval) and ISMER (UQAR) for fish rearing. This work was supported by Ouranos Inc., Ressources Aquatiques Québec (RAQ), and a NSERC strategic grant to LB, DG, and CA (grant number STPGP 521227 - 18).

## Ethics

Animal care was performed humanely under Université Laval’s Comité de protection des animaux permis VRR-18-111 for offspring and Université du Québec à Rimouski’s Comité de protection des animaux permis CPA-76-19-205 for parents.

## Data, code, and materials

Raw Methyl-Seq data and whole genome pool-seq data are being uploaded to NCBI SRA. Analysis codes are available on GitHub as indicated in the methods.

