## Supplementary files for "Thermal regime during parental sexual maturation, but not during offspring rearing, modulates DNA methylation in brook charr (*Salvelinus fontinalis*)"

Figure S1: Brook charr breeding design. Adults were split between warm and cold temperatures during sexual maturation, mated in four modified 2x2 crosses, and the offspring were split between warm and cold temperatures (5 and 8°C) during rearing. Families marked with an asterisk had only one male offspring successfully assigned parentage and thus are not replicated for that experimental group.

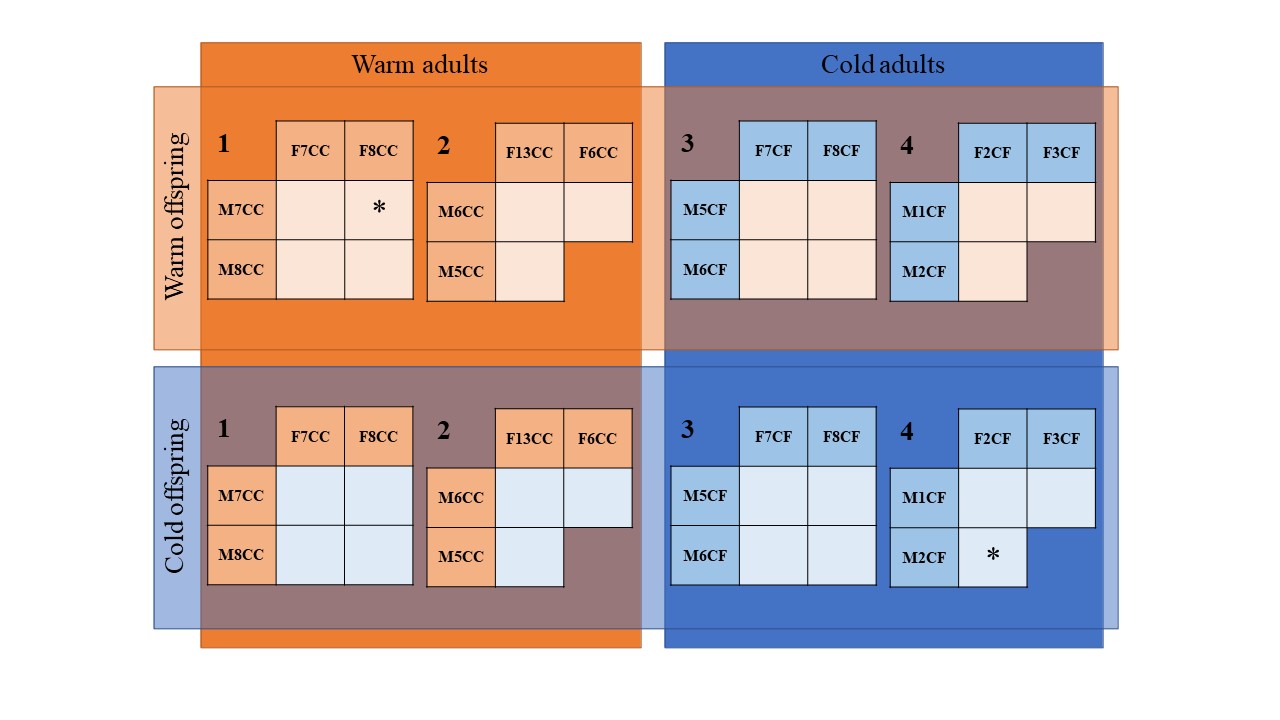

Figure S2: Adult sexual maturation temperature resulted in 464 DMRs at p<0.001 in offspring liver before jackknifing the DSS analysis. Hierarchical clustering along the x-axis based on Euclidean distance for methylation levels shows fairly consistent grouping of full-sibling families, and some evidence of grouping for maternal and paternal half-sibling families.

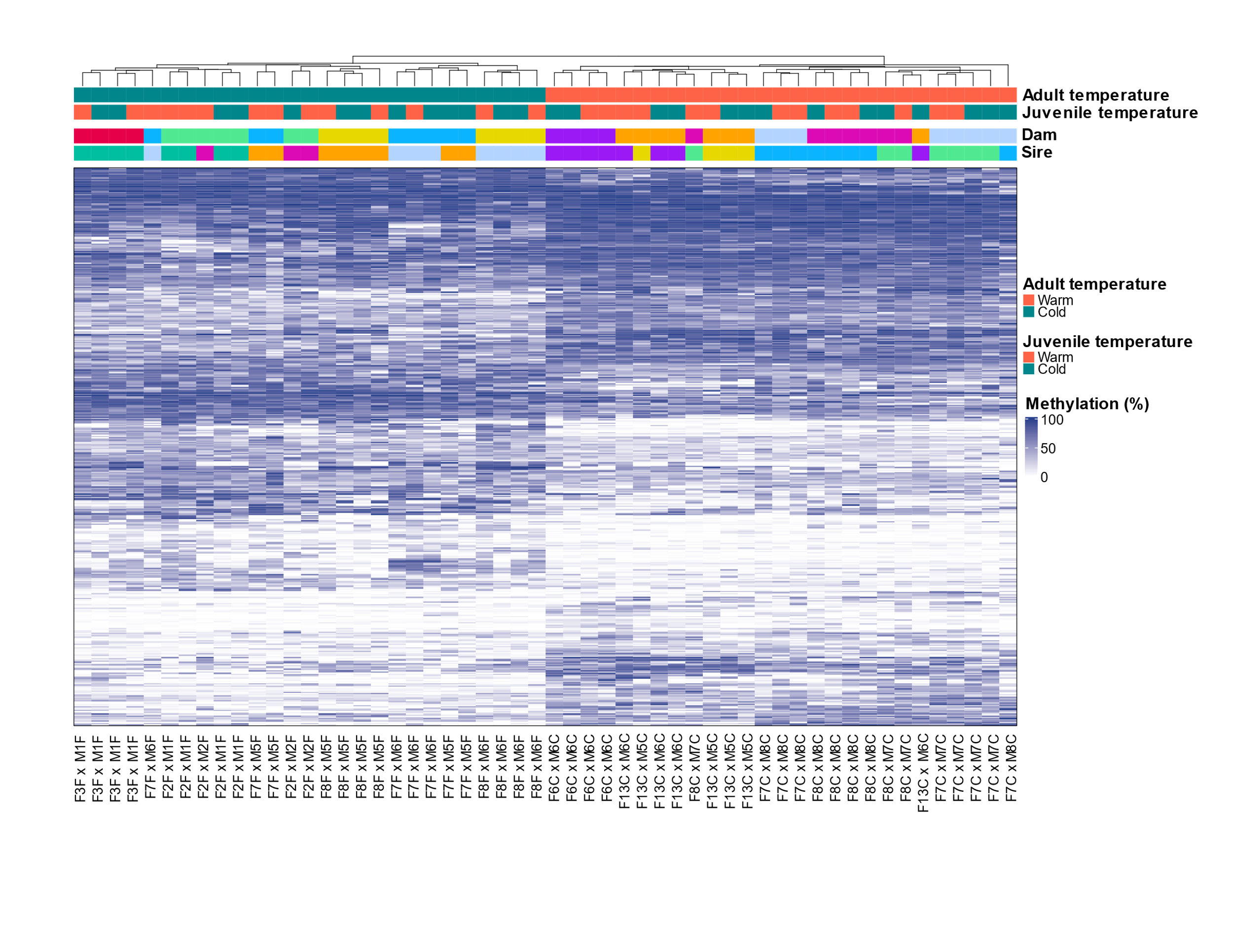

Table S1: GO terms associated with transcripts overlapping DMRs. The number of overlaps indicates the number of unique transcripts directly overlapping jackknifed DMRs. No GO terms showed significant enrichment after FDR correction.

| # Overlaps | GO ID | Term | Ontology | Parent term |
| --- | --- | --- | --- | --- |
| 5 | GO:0007165 | signal transduction | BP |  |
| 4 | GO:0006357 | regulation of transcription by RNA polymerase II | BP |  |
| 4 | GO:0035556 | intracellular signal transduction | BP | GO:0007165 |
| 3 | GO:0000122 | negative regulation of transcription by RNA polymerase II | BP | GO:0006357 |
| 3 | GO:0001525 | angiogenesis | BP |  |
| 3 | GO:0001701 | in utero embryonic development | BP |  |
| 3 | GO:0007049 | cell cycle | BP |  |
| 3 | GO:0007155 | cell adhesion | BP |  |
| 3 | GO:0007264 | small GTPase mediated signal transduction | BP | GO:0007165 |
| 3 | GO:0007420 | brain development | BP |  |
| 3 | GO:0008285 | negative regulation of cell population proliferation | BP |  |
| 3 | GO:0030154 | cell differentiation | BP |  |
| 3 | GO:0045892 | negative regulation of transcription, DNA-templated | BP |  |
| 8 | GO:0046872 | metal ion binding | MF |  |
| 5 | GO:0042802 | identical protein binding | MF |  |
| 4 | GO:0000978 | RNA polymerase II cis-regulatory region sequence-specific DNA binding | MF |  |
| 4 | GO:0000981 | DNA-binding transcription factor activity, RNA polymerase II-specific | MF |  |
| 4 | GO:0003677 | DNA binding | MF |  |
| 4 | GO:0005085 | guanyl-nucleotide exchange factor activity | MF |  |
| 3 | GO:0008270 | zinc ion binding | MF | GO:0046872 |
| 3 | GO:0017124 | SH3 domain binding | MF |  |
| 3 | GO:0051015 | actin filament binding | MF |  |
